# Fas-threshold signalling in MSCs causes tumour progression and metastasis

**DOI:** 10.1101/2020.12.02.406918

**Authors:** Andrea Mohr, Chu Tianyuan, Christopher T. Clarkson, Greg N. Brooke, Vladimir B. Teif, Ralf M. Zwacka

## Abstract

Mesenchymal stem cells (MSCs) are part of the tumour microenvironment and have been implicated in tumour progression. We found the number of MSCs significantly increased in tumour-burdened mice driven by Fas-threshold signalling. Consequently, MSCs lacking Fas lost their ability to induce metastasis development in a pancreatic cancer model. Mixing of MSCs with pancreatic cancer cells led to sustained production of the pro-metastatic cytokines CCL2 and IL6 by the stem cells. The levels of these cytokines depended on the number of MSCs, linking Fas-mediated MSC-proliferation to their capacity to promote tumour progression. Furthermore, we discovered that CCL2 and IL6 were induced by pancreatic cancer cell-derived IL1. Analysis of patient transcriptomic data revealed that high FasL expression correlates with high levels of MSC markers as well as increased IL6 and CCL2 in pancreatic tumours. Moreover, both FasL and CCL2 are linked to elevated levels of markers specific for monocytes known to possess further pro-metastatic activities. These results confirm our experimental findings of a FasL-MSC-IL1-CCL2/IL6 axis in pancreatic cancer and highlight the role MSCs play in tumour progression.

## Introduction

FasL has recently been shown, to induce proliferation of human mesenchymal stem cells (MSCs) *in vitro* (Rippo *et al*, 2013). Although MSCs were found to proliferate in the tumour microenvironment (TME) (Studeny *et al*, 2002), a direct connection between increased MSC numbers and FasL-induced proliferation has so far not been established *in-vivo*. Furthermore, it is not clear whether such increase in MSCs might be linked to their pro-metastatic activity. FasL is commonly known for its pro-apoptotic function via FADD and caspase-8 (Scaffidi *et al*, 1998; Scaffidi *et al*, 1999). However, more recently it was shown that FasL can also trigger non-apoptotic/non-canonical pathways (Kober *et al*, 2011; Lee *et al*, 2011; Legembre *et al*, 2004; Roder *et al*, 2011; Siegmund *et al*, 2007; Wajant *et al*, 2003). Several reports have described this as Fas-threshold signalling, in which low concentrations of FasL resulted in survival signalling and growth (Bentele *et al*, 2004; Lavrik *et al*, 2007; Paulsen *et al*, 2011). In clinical samples, FasL has been found at elevated levels in various cancer types (Belt *et al*, 2014; Botti *et al*, 2004; El-Sarha *et al*, 2009; Herrnring *et al*, 2000; Mottolese *et al*, 2005; Mullauer *et al*, 2000; Muschen *et al*, 1999; Ohta *et al*, 2004; Pernick *et al*, 2002; Sjostrom *et al*, 2002; Song *et al*, 2001; Zhang *et al*, 2005), and has been associated with tumour progression (Barnhart *et al*, 2004; Chen *et al*, 2010; Hoogwater *et al*, 2010; Kleber *et al*, 2008; Lin *et al*, 2012; Malleter *et al*, 2013; Peter *et al*, 2015; Teodorczyk *et al*, 2015; Trauzold *et al*, 2005; Zhang *et al*, 2009; Zheng *et al*, 2013). Most of this work, however, focused on Fas/FasL signalling in cancer cells without consideration of the more complex cellular interactions and regulatory circuits between malignant and non-malignant cells, such as MSCs, present in the TME (Shi *et al*, 2017).

MSCs are initially recruited to the tumour from the bone marrow or other tissues during cancer development (Balkwill *et al*, 2012; Mohr & Zwacka, 2018; Pittenger *et al*, 1999), where they can promote cancer cell dissemination and the development of metastatic lesions (Albarenque *et al*, 2011; Karnoub *et al*, 2007). One of the factors involved in this process, identified in breast cancer metastasis, is the Chemokine (C-C motif) ligand 5 (CCL5), also known as RANTES (Regulated on activation, normal T cell expressed and secreted) (Karnoub *et al*., 2007). CCL5 was shown to be induced by cancer cell-derived osteopontin (OPN) in MSCs (Mi *et al*, 2011). However, CCL5 did not seem to regulate the MSC-induced metastasis of all breast cancer models investigated (Karnoub *et al*., 2007), and its role in other cancer types is also not yet clear. Thus, it seems likely that additional, so far unidentified factors and mechanisms, will have critical roles in MSC-driven tumour progression. As pancreatic cancer is characterised by a pronounced stromal desmoplastic reaction, in which tumour-infiltrating MSCs and carcinoma-associated fibroblasts (CAFs) derived from MSCs become a major component of the TME, we focused on this cancer type (Chan *et al*, 2019; Kamisawa *et al*, 2016). Moreover, the majority of cases of pancreatic cancer are complicated by distant metastasis (Kamisawa *et al*, 1995) highlighting the need to better understand the process of tumour progression of this persistently difficult to treat disease. Therefore, we examined the action of FasL on MSCs and the link to pro-metastatic behaviour of MSCs in pancreatic cancer.

## Materials and Methods

### Cell culture and reagents

The colorectal carcinoma cell lines HT29 (ATCC, Manassas, VA, USA) and Colo205 (ATCC) were cultured in RPMI (Lonza, Basel, Switzerland) and in McCoy’s 5A (Lonza), respectively. The prostate cancer cell lines, LNCaP (ATCC) and PC3 (ATCC), were cultured in RPMI medium. The breast cancer cell lines MDA-MB-231 (ATCC), and MCF7 (ATCC) were cultured in DMEM (Lonza) and in EMEM (Lonza) medium, respectively. The pancreatic cancer cells PancTu1 (obtained from Prof Simone Fulda), AsPC1 (ATCC), BxPC3 (ATCC) were cultured in RPMI medium, while MiaPaCa2 (ATCC) cells were grown in DMEM medium. All media were supplemented with 10% foetal bovine serum (FBS) (Thermo Fisher Scientific, Waltham, MA, USA), 100 U/ml penicillin and 100 μg/ml streptomycin.

Human bone marrow derived MSCs were from Lonza, umbilical cord derived MSCs were purchased from Promocell (Heidelberg, Germany), and human adipose derived stem cells were from Amsbio (Cambridge, MA, USA). Human MSCs were cultured in StemMACS MSC Expansion Medium (Miltenyi Biotec, Bergisch Gladbach, Germany). Murine MSCs and LPR-MSCs were isolated and cultured as previously described (Albarenque *et al*., 2011; Drappa *et al*, 1993; Watanabe-Fukunaga *et al*, 1992). The following inhibitors were used: STAT3i (C188-9), PI3Ki (AZD8186 and BYL719), ERKi (PD0325901), p38i (R1503), NF-kBi (Bay11), JNKi (SP600125). All inhibitors were purchased from Stratech (Ely, UK). The following cytokines were ordered from Peprotech (Rocky Hill, NJ, USA): IGF1, IL1α, IL1β, OSM, PDGF and VEGF. TRAIL, IL6, CCL2, anti-CCL2 antibody, human IL6Rα and human gp130 were purchased from Biotechne (Minneapolis, MN, USA), FasL from Enzo (Farmingdale, NY, USA), multimeric FasL and anti-APO-1-3 from AdipoGen Life Sciences (San Diego, CA, USA). IL1RA was purchased from Biolegend (San Diego, CA, USA) and anti-IL6 antibody from Diaclone (Besancon, France).

### Animal studies

Ten-week-old female CD1 nu/nu mice (Charles River) were subcutaneously injected with 5×10^6^ tumour cells (MDA-MB-231 and PancTu1, respectively) in 200 μl PBS under the lateral skin of the right leg. When tumours were palpable, the animals were injected intravenously with 1×10^5^ Chloromethyl-dialkylcarbocyanine (CM-Dil; Thermo Fisher Scientific) labelled MSCs. For the analysis of the tissue distribution of the MSCs and their effects on metastasis formation inguinal, axillary, mesenteric and paraaortic lymph nodes, liver, spleen, kidney, lungs and tumour were fixed in 10% neutral buffered formalin. In addition, tibia and femur, and vertebra of each animal were formalin fixed, followed by decalcification with a solution of 8% formic acid/8% HCl prior to tissue processing and paraffin embedding. For histopathological analyses, 4 μm-paraffin sections were stained with Haematoxylin and Eosin (H&E) and examined using light microscopy. The animal studies were performed according to national laws and covered by license from the Irish government.

To detect MSCs *in vivo* they were loaded with 4 μg/ml CM-Dil in pre-warmed PBS for 15 min at 37° C followed by an incubation for 15 min at 4° C. The cells were washed with PBS, resuspended in their normal growth medium and cultured for 16 h before they were used. For quantification of MSC bio-distribution, 4 μm paraffin sections were deparaffinised and rehydrated, washed in PBS, followed by incubation for 10 min with DAPI, and mounted in fluoromount solution for the detection of fluorescent cells under a microscope. 15 randomly chosen fields/organ of each animal were counted.

### Cell surface staining

1×10^6^ cells were incubated with fluorochrome conjugated antibodies for 15 minutes on ice in the dark. Before flow cytometric analyses, the cells were washed with PBS containing 1% BSA. The following PE-conjugated antibodies were used for cell surface staining: human CD95 (DX2), mouse CD95 (SA367H8). PE-conjugated Mouse IgG1κ was used as isotype control. All antibodies used were purchased from BioLegend and were used at a concentration of 0.2 μg/10^6^ cells.

### Cell growth assays

MSCs were plated at a density of 1×10^4^ in a 24-well plate. After 24 h medium was changed to DMEM containing 1% FCS. 12 h later, the cells were treated with 0.2 ng/ml FasL, 0.2 ng/ml multimeric FasL, and 0.2 ng/ml anti-APO-1-3, respectively, or at concentrations indicated in the figure legends. As a control, cells were treated with 0.2 ng/ml TRAIL. JNKi, p38i, ERKi were added at a concentration of 1 µM, 20 min before FasL was added. To measure cell numbers, cells were detached with trypsin solution and counted using a haemocytometer. Crystal violet staining was performed after 7 days as previously described (Mohr *et al*, 2019).

### Apoptosis assays

MSCs were plated at a density of 5×10^4^ in a 24-well plate. After 24 h, cells were treated with FasL, multimeric FasL, or anti-APO-1-3. Apoptosis was measured according to Nicoletti et al. (Nicoletti *et al*, 1991). Briefly, 24 h post-treatment, cells were harvested and washed once with PBS. Cells were resuspended in hypotonic fluorochrome solution containing 50 g/ml propidium iodide, 0.1% sodium citrate, and 0.1% Triton-X100. After incubation at 4° C for 2 h, cells were analysed by flow cytometry.

### Western blot

MSCs were serum starved for 24 h, before 0.2 ng/ml FasL was added. Cells were harvested at the indicated time points. For analysing the effects of tumour cell supernatants on MSCs, the stem cells were seeded and 24 h later their growth medium was replaced with cell supernatant for the indicated time-points. Protein transfers onto PVDF membranes were performed as described previously (Mohr *et al*, 2018). The following primary antibodies were used: anti-phospho-SAPK/JNK (Cell Signaling Technology, Danvers, MA, USA), anti-total-SAPK/JNK (Cell Signaling Technology), anti-phospho-p38 MAPK (Cell Signaling Technology), anti-total-p38 MAPK (Cell Signaling Technology), anti-phospho-ERK (Cell Signaling Technology), anti-total-ERK (Cell Signaling Technology), anti-CD95 (Santa Cruz Biotechnology, Dallas, TX, USA) and anti-IκBα (Cell Signaling Technology), and anti-CuZnSOD (Binding Site, Birmingham, UK). Peroxidase-conjugated secondary antibodies were anti-rabbit and anti-goat from Santa Cruz. All antibodies were diluted to a working concentration of 1:1000.

### NF-κB reporter assay

MSCs were seeded in 12-well plates. Next day, cells were adenovirally transduced with an NF-κB-firefly-luciferase vector (Vector Biolabs, Malvern, PA, USA) at an MOI of 200. 24 h after transduction, the cells were pre-treated with 1 μM NF-κB inhibitor or IL1RA (4 μg/ml) for 20 min before tumour cell supernatant was added. 24 h post treatment, cells were lysed in passive lysis buffer (Promega, Madison, WI, USA). Luciferase activity was measured in a luminometer.

### ELISA

1×10^4^ tumour cells were either mixed with varying numbers of MSCs (1×10^4^ MSCs (1:1), 1×10^3^ MSCs (1:10)) or treated with tumour cell supernatants for 2 days. To test for cytokine induction, 1×10^4^ cells were cultured and subsequently treated with 0.2 ng/ml of the respective death ligands, 50 ng/ml of IGF1, IL1α, IL1β, OSM, PDGF-BB, TPO or VEGF-A, 2.5 μg/ml shIL6Rα, 2.5 μg/ml IgG1κ, 2.5 μg/ml anti-IL6, 2.5 μM STAT3i, different PI3K inhibitors (PI3Ki-1: AZD8186, PI3Ki-2: BYL719) and the NK-κB inhibitor at concentrations of 1 μM, 4 μg/ml IL1RA or 2.5 μg/ml gp130-Fc for 48 h. Supernatants were removed and cleared by centrifugation. The human and murine cytokines IL6, CCL2 and CCL5 levels were determined by DuoSet ELISA (Biotechne), according to manufacturer’s instructions.

For the analysis of sustained secretion of CCL2 and IL6 by MSCs, supernatants from PancTu1 cells were applied to 1×10^4^ MSCs. The medium was changed after 3 days to fresh medium and replaced after another 3 days. Cells without medium change served as control. 50 μl medium samples were taken every 3 days. The 50 μl taken for experiments were replaced with fresh medium. For the measurement of sIL6R, IL1RA, IL1α and IL1β, the supernatant was taken from cells cultured for 3 days. The amount of supernatants used for these ELISAs were adjusted to the respective cell number.

### Cytokine Array

For these studies, we took supernatants from hMSCs and tumour cells at a ratio of 1 MSC:10 tumour cells and compared the cytokine levels to MSCs and tumour cells cultured separately. The supernatants were diluted 3.5-fold and applied to human cytokine antibody arrays III (RayBiotech, Norcross, GA, USA) according to manufacturer’s instructions. We normalised each cytokine signal to its internal control and calculated the fold upregulation.

### Invasion assay

Tumour cells were labelled with CM-Dil and serum starved for 24 h. The labelled cells were then cultured alone, treated with 50 ng/ml cytokines or mixed with MSCs in the upper part of Matrigel coated Boyden chambers (8 μM, BD Biosciences, San Jose, CA, USA). Cell invasion was determined by analysing the number of labelled cells that migrated through the Matrigel. The effect of CCL2 or IL6 was determined by adding 2 μg/ml anti-CCL2 antibody, 10 μg/ml anti-IL6 antibody or 4 μg/ml IL1RA to the tumour cell/MSC mixtures. The control was mouse IgG2b.

### Differentiation of promyelocytic cells to macrophage-like cells

HL60 cells were plated at a density of 4×10^5^ cells/ml and differentiated with 12-O-tetradecanoylphorbol-13-acetate (TPA) (Sigma, Darmstadt, Germany) at a final concentration of 32 nM for 6 days. For the determination of CCL2, differentiated and undifferentiated cells were added to MSCs treated with conditioned medium from PancTu1 cells for 48 h.

### TCGA expression analysis

Expression data from the pancreatic cancer patient cohort were downloaded from the Cancer Genome Atlas (TCGA) Research Network and can be obtained at https://cancergenome.nih.gov/. RNaseq V2 level 3 data were downloaded for 186 pancreatic cancer samples from the TCGA data portal. The z-score threshold was set at 1.5. The downloaded dataset was parsed and visualized in R. The heatmaps were plotted using the R package ComplexHeatmap (Gu *et al*, 2016). The script used is available at https://github.com/chrisclarkson/FASLG_heatmap_analysis.

## Results

### Tumour-burdened animals show an increase in MSC numbers

In order to study the fate of murine (mMSCs) and human (hMSCs) MSCs in tumour-burdened animals we initially used the MDA-MB-231 mammary carcinoma model, for which MSC-promoted metastasis development had been shown before (Albarenque *et al*., 2011; Karnoub *et al*., 2007). To analyse the biodistribution of MSCs, we intravenously injected CM-Dil labelled MSCs when the MDA-MB-231 xenografts became palpable (Figure 1A). For both mMSCs (Figure 1B) and hMSCs (Figure 1C) we found an increase in MSC numbers in the tumour-burdened animals as compared to those carrying no xenograft. This increase only became visible four-weeks post-injection, whereas it was not apparent after one week, demonstrating that tumour-induced MSC proliferation is the cause of the observed effect.

**Figure 1.**
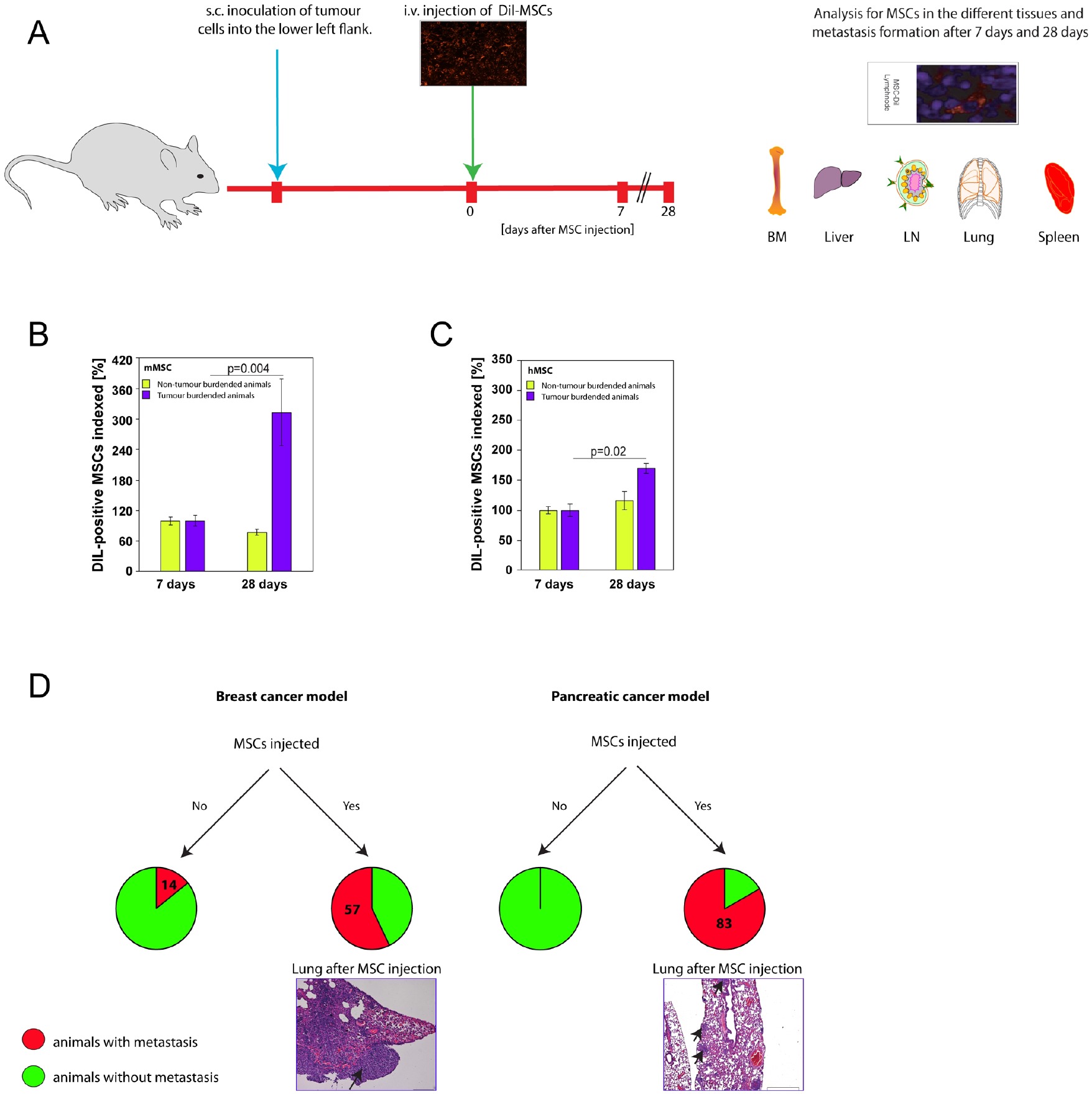
Tumour-burdened animals show an increase in MSC numbers. **A**. Schematic overview of experimental design to study MSC biodistribution, numbers and metastasis development. **B, C**. Relative numbers of murine MSCs (mMSCs) and human MSCs (hMSCs) found in control (yellow) and tumour-burdened (MDA-MB-231) animals (purple) after 7 days and 28 days, respectively. The relative numbers of MSCs found in the following tissues are shown in the diagram: bone marrow, lymph nodes, liver, lung and spleen. The average numbers on day 7 were set to 100% to allow for comparison to the 28-day time point. Each cohort consisted of three animals. **D**. Incidence (%) of metastases in different tumour models (breast cancer, left; pancreatic cancer, right) after systemic murine MSC administration. Representative images of H/E stained lung microsection with metastatic lesions (black arrows) from tumour-burdened mice injected with MSCs are also shown. Each cohort consisted of six animals.

In order to study the impact of this effect on tumour progression we used, in addition to the MDA-MB-231 breast cancer cells, a PancTu1 pancreatic cancer model (Figure 1A). While MDA-MB-231 cells show a degree of metastasis without MSCs, PancTu1 cells do not when grown as subcutaneous xenograft (Alves *et al*, 2001). Therefore, we examined first whether MSCs were able to induce metastasis development in the PancTu1 model. Four weeks after the injection of MSCs we examined the lungs from animals with MDA-MB-231 and PancTu1 xenografts for metastasis formation on H&E stained sections. In the cohorts that received MSCs, we found more than half of the mice had developed metastatic lesions in the MDA-MB-231 model, and 83% in the mice with PancTu1-derived xenografts (Figure 1D). Contrary, in animals that did not receive MSCs we could detect lung metastatic nodules in only 14% for MDA-MB-231 tumours, and as expected, none for PancTu1 xenografts (Figure 1D). These results demonstrate that MSCs can exert their pro-metastatic function onto cells that normally do not disseminate providing a cleaner and better model to study the effect of tumour-induced MSC proliferation on metastasis development.

### FasL induces proliferation in MSCs via MAPK/ERK

Increased FasL levels have been detected in various cancer types including breast and pancreatic cancer (El-Sarha *et al*., 2009; Mullauer *et al*., 2000; Muschen *et al*., 1999; Ohta *et al*., 2004; Pernick *et al*., 2002). This can provide the proliferative signal for circulating MSCs and/or MSCs in the TME. However, FasL is a member of the TNF superfamily (TNFSF) and a so-called death ligand that is normally associated with apoptosis induction. In order to reconcile the pro-apoptotic function with a role in MSC proliferation and consequent tumour progression, MSCs would have to be resistant to FasL-induced apoptosis. To test whether MSCs are principally able to respond to FasL we measured the levels of its receptor, Fas, on the surface of different types of hMSCs and mMSCs. While we found robust expression for all MSC types demonstrating that they should be able to respond (Figure 2A), all of them were resistant to up to 5 ng/ml of FasL (Figure 2B). Treatment with agonistic Fas antibodies also failed to trigger cell death in MSCs (Supplementary Figure 1A). Only when we used potent multimeric-FasL at high concentrations (5 ng/ml) we could detect apoptosis in both hMSCs and mMSCs (Supplementary Figure 1A). These findings indicate that MSCs have the potential to trigger cell death, but that Fas is configured to respond very weakly to its ligand with regard to apoptosis-induction. In contrast, treatment of different types of MSCs with low levels of FasL (0.2 ng/ml) gave rise to increased proliferation in human and murine MSCs over a period of five days (Figure 2C). Similar results were achieved in an outgrowth assay (Supplementary Figure 1B). At higher concentrations of FasL the proliferative activity was lost and MSCs no longer grew more than controls over the five-day period (Figure 2D). In addition, we examined whether agonistic Fas antibodies could also induce proliferation in MSCs. We found that in contrast to FasL and multimeric-FasL, treatment with these antibodies did not give rise to enhanced proliferation in MSCs (Supplementary Figure 1C). As FasL acts as trimers engaging trimeric Fas receptor complexes, whereas agonistic antibodies tend to lead to dimeric forms, we concluded that receptor trimers are required for Fas signaling and proliferation of MSCs. In contrast to the findings with FasL, TRAIL, another apoptosis-inducing member of the TNFSF, did not lead to increased proliferation of MSCs (Supplementary figure 1D).

**Figure 2.**
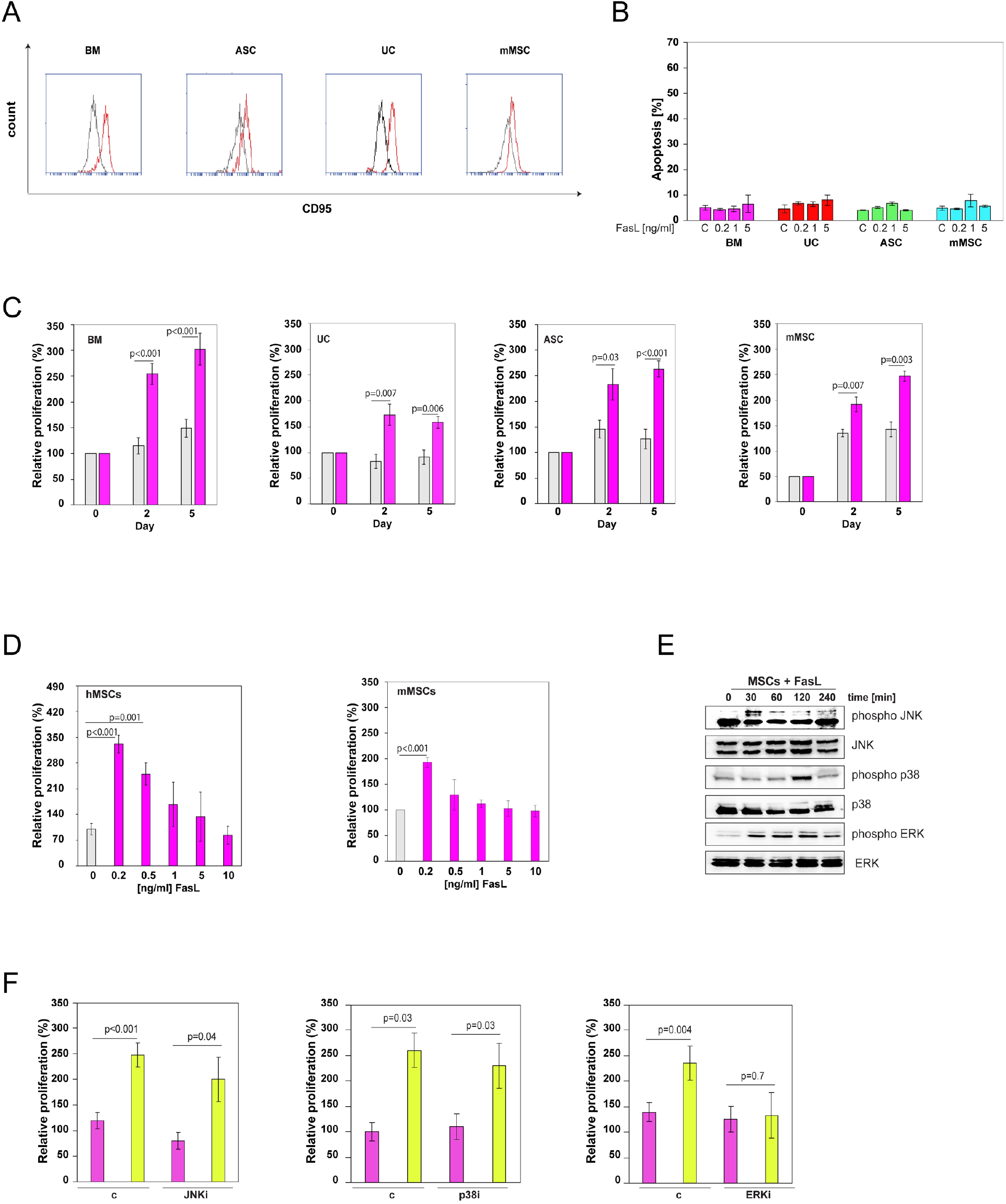
FasL induces proliferation in MSCs via MAPK/ERK. **A**. Representative histograms of flow cytometric analyses of Fas surface expression in human MSCs from bone marrow (BM), adipose tissue (ASC) and umbilical cord (UC) as well as murine MSCs (mMSC). **B**. Apoptosis measurements in different types of MSCs after treatment with recombinant FasL for 24 h, at concentrations ranging from 0.2-5 ng/ml. Controls (c) were treated with carrier only. Results are the mean ± SEM; n=3. **C**. Human MSCs from bone marrow (BM), umbilical cord (UC) and adipose tissue (ASC) as well as murine MSCs (mMSC) were treated with 0.2 ng/ml of FasL for 2 and 5 days and cell numbers determined (purple). MSCs treated with carrier are shown as controls (grey). The numbers of MSCs at the start of FasL treatment (Day 0) were set to 100%. Results are the mean ± SEM; (BM-MSCs ASC-MSCs n=8; UC-MSCs n=5; mMSCs n=6). **D**. Human (hMSC, left) and murine MSCs (mMSC, right) were treated with different concentrations of FasL, as shown in the figure, for 2 days. The numbers of MSCs at the start of FasL treatment (Day 0) were set to 100%. Results are the mean ± SEM; n=8. **E**. Murine MSCs were treated with 0.2 ng/ml FasL for 30, 60, 120 and 240 min and the resulting protein lysates analysed by western blot with phospho-specific antibodies against JNK, p38 and ERK. Western blots with regular antibodies served as loading controls. **F**. Murine MSCs were treated with 0.2 ng/ml FasL for 2 days, and JNKi (left), p38i (centre) and ERKi (right), respectively. The numbers of MSCs at the start of FasL treatment (Day 0) were set to 100%. Results are the mean ± SEM; n=4. Carrier treated controls (c) are shown as comparison.

Next, we analysed a number of non-apoptotic pathways that can be triggered by Fas signalling. Treatment of MSCs with 0.2 ng/ml FasL, and subsequent western blotting of the resulting protein lysates, showed that MAPK/ERK was substantially phosphorylated/activated, whereas JNK and p38 only showed a very transient peak in phosphorylation/activation (Figure 2E). The use of inhibitors revealed that blocking JNK or p38 had no impact on FasL-induced MSC proliferation (Figure 2F). In contrast, MAPK/ERK inhibitors reduced proliferation back to normal control levels (Figure 2F). Thus, FasL acts via MAPK/ERK to induce proliferation in MSCs.

### Pancreatic cancer cells stimulate MSCs to produce the pro-metastatic cytokines CCL2 and IL6

The chemokine CCL5 and its receptor CCR5 have been implicated in MSC-mediated breast cancer metastasis (Karnoub *et al*., 2007). Breast cancer cells were shown to stimulate secretion of CCL5 from MSCs, which then acts in a paracrine fashion on the cancer cells to enhance their motility, invasion and metastasis. Therefore, we examined first, whether FasL could directly induce CCL5 expression in MSCs, while at the same time stimulating their proliferation. When we added FasL (or multimeric FasL; or agonistic Fas antibodies) to human or murine MSCs we could not detect an increase in the levels of CCL5 by ELISA (Figure 3A). Next, we tested whether cross-talk with pancreatic cancer cells led to CCL5 induction in MSCs, similar to MDA-MB-231 breast cancer cells. Indeed, when we mixed MDA-MB-231 cells with increasing numbers of mMSC, we could measure substantial CCL5 production and secretion in an MSC-dose dependent manner (Figure 3B). However, when we used PancTu1 pancreatic cancer cells in such mixing experiments, we could not detect CCL5 induction irrespective of the number of MSCs (Figure 3B). The same pattern of CCL5 production was seen with hMSCs, where MDA-MB-231 cells led to induction of CCL5 but PancTu1 cells did not (Figure 3C). Testing three additional pancreatic cancer cell lines (MiaPaCa2, AsPC1 and BxPC3) resulted in the same outcome (Figure 3D). Thus, CCL5 cannot explain the MSC-induced metastasis development in our pancreatic cancer model (Figure 1D). Therefore, we screened mixes of PancTu1 cells and hMSCs on a cyto-/chemokine array for factors that were induced by the interaction of the two cell types. We discovered that CCL2 was upregulated and secreted into the supernatant medium (Figure 3E). In addition, we could also detect significant increases of IL6 production, albeit starting from a higher basal level. Other factors on the array such as IL8, GRO chemokines (CXCL1, CXCL2 and CXCL3) and several others were not affected (Figure 3F). The induction of CCL2 and IL6 were corroborated by ELISA, which showed rising levels of the two cytokines with increasing MSC numbers, i.e. mixing more MSCs with the same number of PancTu1 cells led to more cytokine production (Figure 3G). Moreover, mMSCs responded in the same way to mixing them with pancreatic cancer cells (Figure 3H), These results point to MSCs as a specific cellular production site for CCL2 and IL6 in response to interaction with pancreatic cancer cells. This hypothesis was confirmed by murine CCL2-ELISAs of supernatants of PancTu1/mMSCs and PancTu1/hMSCs mixes, of which only the former provided a positive result (Figure 3H).

**Figure 3.**
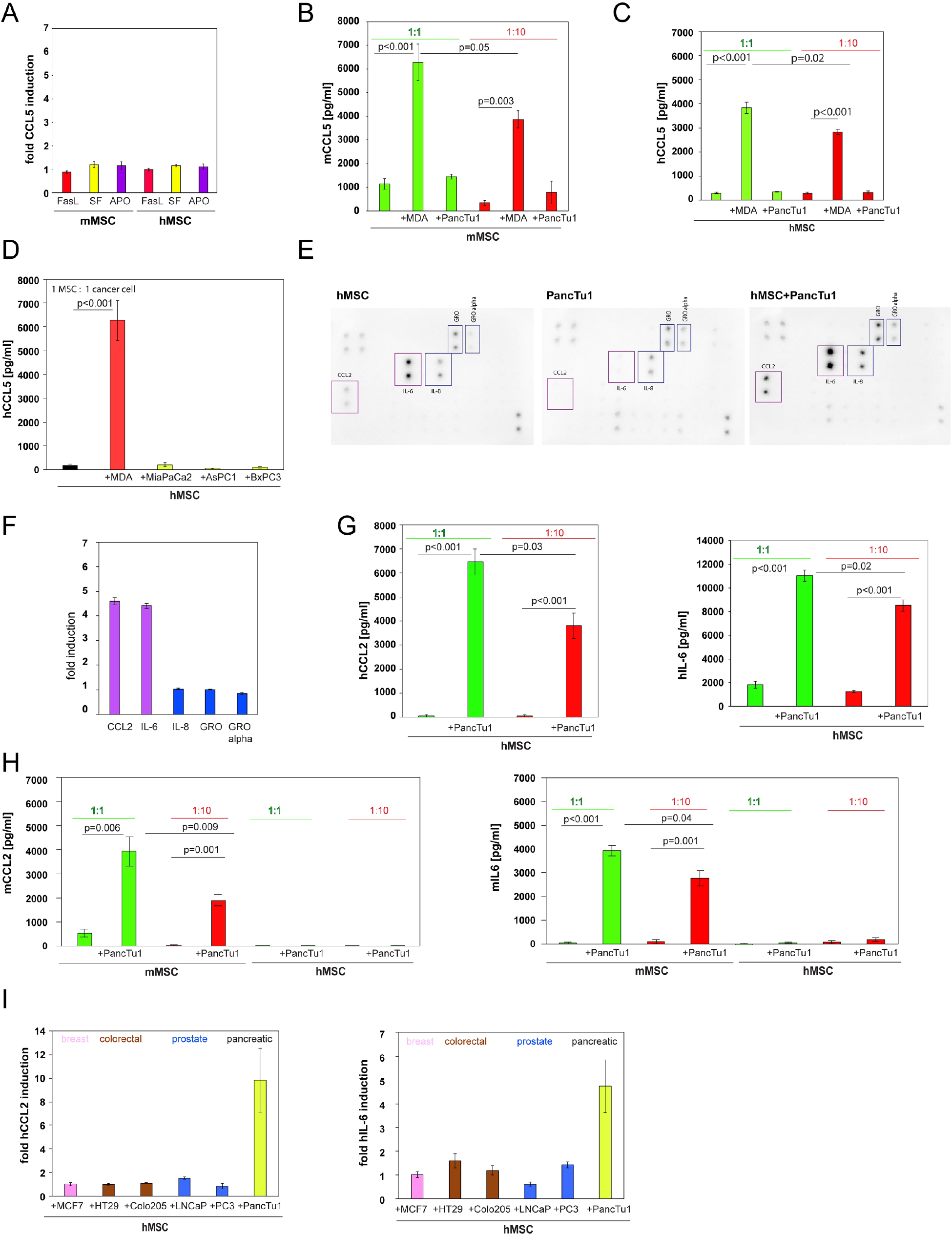
Pancreatic cancer cells stimulate MSCs to produce the pro-metastatic cytokines CCL2 and IL6. **A**. Murine (mMSC) and human MSCs (hMSC) were treated with 0.2 ng/ml FasL, 0.2 ng/ml multimeric FasL (SF) and 0.2 ng/ml of an anti-APO-1-3 antibody (APO) for 72 h. The resulting supernatants were analysed for CCL5 by ELISA. Results are the mean ± SEM; n=3. **B**. Murine MSCs (mMSCs) were mixed with MDA-MB-231 (MDA) and PancTu1 cells at ratios of 1:1 and 1:10. The resulting supernatants were analysed for CCL5 by ELISA. Results are the mean ± SEM; n=3. **C**. Human MSCs (hMSCs) were mixed with MDA-MB-231 (MDA) and PancTu1 cells at ratios of 1:1 and 1:10. The resulting supernatants were analysed for CCL5 by ELISA. Results are the mean ± SEM; n=3. **D**. Human MSCs (hMSCs) were mixed with three different pancreatic cancer cell lines (MiaPaCa2, AsPC1 and BxPC3) at a ratio of 1:1 and the resulting supernatants analysed for CCL5 by ELISA. Mixing experiments with MDA-MB-231 (MDA) cells were carried out as positive control. Results are the mean ± SEM; n=3. **E**. Human MSCs (hMSCs) were mixed with PancTu1 cells at a ratio of 1:10 for 48 h and the supernatant applied to a cytokine antibody array (right). Supernatants from hMSCs (left) and PancTu1 cells (centre), respectively, were used as controls. **F**. Quantification of selected cytokines (boxed-in in Figure 3E) from the cytokine antibody array. **G**. Human MSCs (hMSCs) were mixed with PancTu1 cells at ratios of 1:1 and 1:10. The resulting supernatants were analysed for CCL2 (left) and IL6 (right) by ELISA. Results are the mean ± SEM; n=3. **H**. Murine MSCs (mMSCs) and human MSCs (hMSCs) were mixed with PancTu1 cells at ratios of 1:1 and 1:10. The resulting supernatants were analysed for CCL2 (left) and IL6 (right) using murine-specific ELISAs. Results are the mean ± SEM; n=3. **I**. Human MSCs (hMSCs) were mixed with different cancer cell lines (MCF7, HT29, Colo205, LNCaP, PC3) at a ratio of 1:1. The resulting supernatants were analysed for CCL2 (left) and IL6 (right) by ELISA. PancTu1 cells were used as positive controls. Results are the mean ± SEM; n=3 (CCL2); n=4 (IL6).

When we used cells from other cancer types (breast, colorectal and prostate cancer) and mixed them with hMSCs we could not detect CCL2 and IL6 production (Figure 3I) pointing to a specific effect in pancreatic cancer.

### A pancreatic cancer cell-derived soluble factor is responsible for the induction of CCL2 and IL6 in MSCs

Pro-metastatic functions have been attributed to CCL2 and IL6 in various cancer types. Therefore, we analysed the direct action of recombinant CCL2 and IL6 on pancreatic cancer cells. In Matrigel assays we found that recombinant CCL2 increased the invasiveness of PancTu1 cells, whereas IL6 showed only a moderate and non-significant effect (Figure 4A). Using PancTu1/MSC mixes plus neutralising CCL2 or IL6 antibodies revealed that only blocking of CCL2 significantly reduced the invasiveness of the PancTu1 cells (Figure 4B). These results demonstrate that pancreatic cancer cells can directly respond to MSC-produced CCL2.

**Figure 4.**
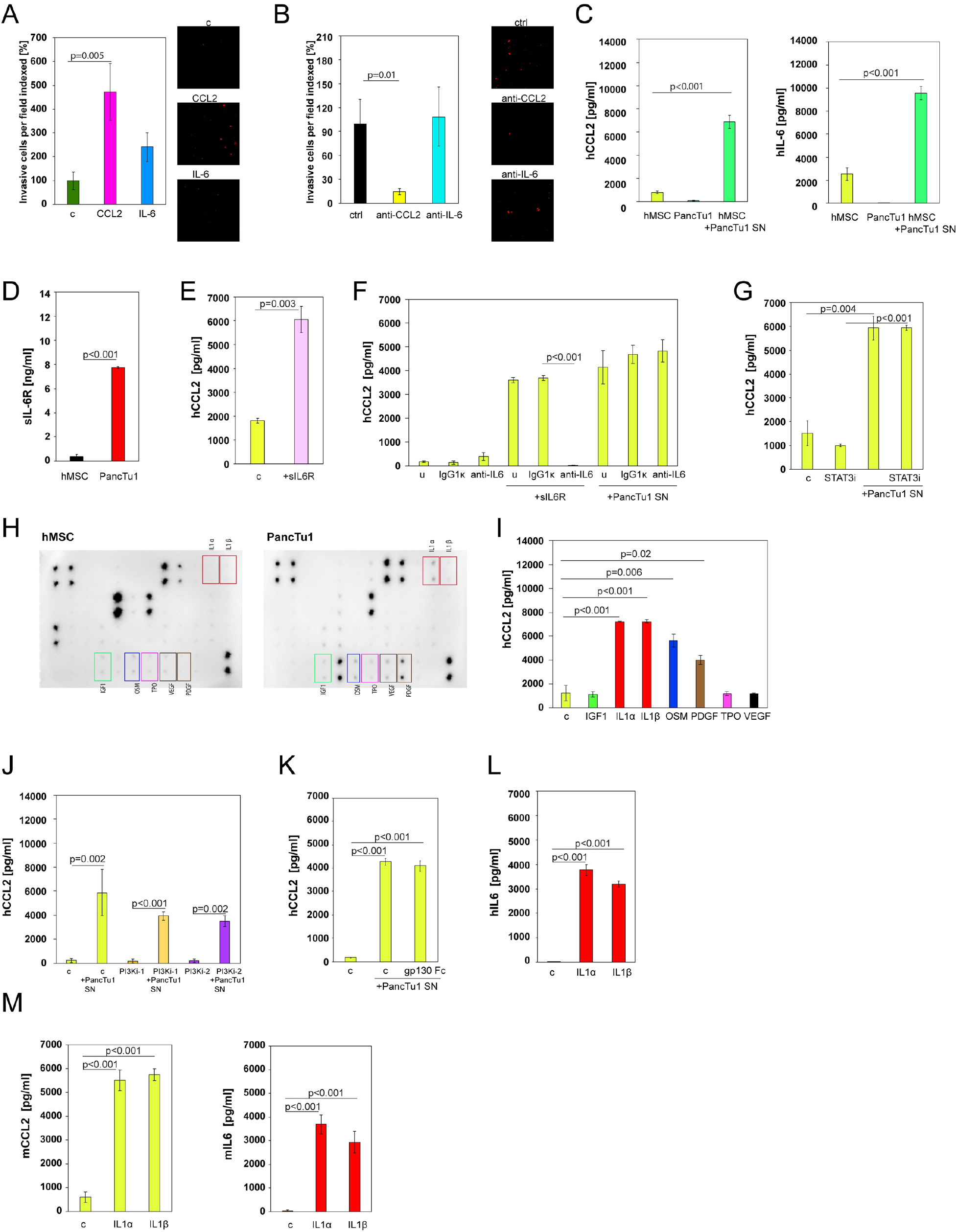
A pancreatic cancer cell-derived soluble factor is responsible for the induction of CCL2 and IL6 in MSCs. **A**. CM-Dil labelled PancTu1 cells were cultured in the presence of carrier (c), CCL2 or IL6 in the upper well of Boyden chambers. Representative images (right) show the labelled PancTu1 cells that migrated through Matrigel coated membranes in 72 h. Quantification of the migration activity of the cells is shown on the left. Results are the mean ± SEM of 5 randomly chosen fields per experiment; n=3. **B**. CM-Dil labelled PancTu1 cells were cultured in the presence of human MSCs with anti-CCL2, anti-IL6 or a control (ctrl) antibody in the upper wells of Boyden chambers. Representative images (right) show the labelled PancTu1 cells that migrated through Matrigel coated membranes in 72 h. Quantification of the migration activity of the cells is shown on the left. Results are the mean ± SEM of 5 randomly chosen fields per experiment; n=2. **C**. CCL2 (left) and IL6 (right) ELISAs of supernatants from human MSCs (hMSCs) that were treated with conditioned medium of PancTu1 cells (SN) for 72 h. Supernatants of PancTu1 cells and MSCs are shown as controls. Results are the mean ± SEM; n=3. **D**. ELISA for sIL6R of supernatants from human MSCs (hMSCs) and PancTu1 cells. Results are the mean ± SEM; n=3. **E**. CCL2 ELISA of human MSC supernatants after treatment with recombinant sIL6R for 72 h. Carrier controls (c) are depicted as controls. Results are the mean ± SEM; n=3. **F**. CCL2 ELISA of supernatants from human MSCs treated with conditioned medium of PancTu1 cells (SN) supplemented with neutralising antibodies against IL6. IgG1κ antibodies and MSCs treated with recombinant sIL6R were used as controls. The results of untreated samples (u) are also shown. Results are the mean ± SEM; n=3. **G**. CCL2 ELISA of supernatants from human MSCs treated with conditioned medium of PancTu1 cells (SN) and a STAT3 inhibitor (STATi). Carrier-treated MSCs (c) and MSCs treated with STAT3 inhibitor alone are shown as controls. Results are the mean ± SEM; n=3. **H**. Long exposure images of cytokine antibody arrays probed with supernatants of human MSCs (hMSCs) and PancTu1 cells, respectively. Differentially expressed growth factors and cytokines are highlighted. **I**. CCL2 ELISA of supernatants from human MSCs treated with different recombinant cytokines (IGF1, IL1α, IL1β, OSM, PDGF, TPO and VEGF). Carrier-treated controls (c) are also shown. Results are the mean ± SEM; n=3. **J**. CCL2 ELISA of supernatants from human MSCs treated with conditioned medium from PancTu1 cells (SN) in the presence of two different PI3K inhibitors. Carrier-treated MSCs (c) and inhibitors alone were used as controls. Results are the mean ± SEM; n=3. **K**. CCL2 ELISA of supernatants from human MSCs treated with conditioned medium from PancTu1 cells (SN) and gp130-Fc. Carrier-treated MSCs (c) are shown as controls. Results are the mean ± SEM; n=3. **L**. IL6 ELISA of supernatants from human MSCs treated with IL1α and IL1β carrier control (c) is also depicted in the graph. Results are the mean ± SEM; n=3. **M**. CCL2 (left) and IL6 (right) ELISAs of supernatants from murine MSCs treated with IL1α and IL1β Carrier controls (c) are also depicted in the graphs. Results are the mean ± SEM; n=3.

In order to understand the mechanisms behind the up-regulation of CCL2 and IL6 in MSCs, we investigated whether cell-to-cell interactions are necessary for induction of these cytokines. Transferring conditioned medium from PancTu1 cells onto hMSCs revealed increased levels of CCL2 and IL6 after 72 h (Figure 4C). The same results were obtained with mMSCs (Supplementary Figure 2). These results again confirm that MSCs produce the two cytokines and show that a soluble factor secreted by the pancreatic cancer cells is responsible for the CCL2 and IL6 induction. Interestingly, hMSCs express relatively high basal levels of IL6 (Figure 3E), which is known to be able to stimulate CCL2 as well as its own expression (Hurst *et al*, 2001). Therefore, we hypothesised that IL-6R trans-signalling (Schaper & Rose-John, 2015), in which constitutive IL6 secreted by MSCs acting together with cancer cell-derived sIL6R, causes the production of CCL2 and additional IL6. Examining the supernatant of PancTu1 cells indeed revealed that they produce sIL6R, albeit at low levels (Figure 4D). In addition, treatment of hMSCs with recombinant sIL6R enhanced CCL2 production (Figure 4E). However, when we used IL6 neutralising antibodies with PancTu1 supernatants on MSCs the CCL2 induction was not reduced (Figure 4F). We observed the same outcome with a STAT3 inhibitor confirming that IL6 (trans-) signalling was not responsible for the effects exerted by the pancreatic cancer cells (Figure 4G). This led us to examine the MSC and PancTu1 cytokine arrays using longer exposure images, this time not focusing on induced factors, but those that are present in the supernatants of the cancer cells but lacking from the medium of MSCs. We identified a total of seven cytokines and growth factors, IGF1, IL1α, IL1β, OSM, PDGF-BB, TPO and VEGF-A (Figure 4H). Of these, recombinant IL1α, IL1β, OSM and PDGF-BB were able to induce CCL2 expression in hMSCs (Figure 4I). As neither a PI3K inhibitor to block PDGF signaling nor recombinant gp130 to block OSM could inhibit the CCL2 induction by PancTu1 medium supernatants, IL1 (IL1α and IL1β) was left as the remaining candidate (Figure 4J and 4K). A role for IL1α and IL1β was supported by the observation that they could also induce IL6 in human MSCs (Figure 4L) as well as CCL2 and IL6 in mMSCs (Figure 4M).

### Pancreatic cancer-derived IL1 induces CCL2 and IL6 via NF-κB activation in MSCs

In order to demonstrate the causative role of IL1 in the supernatants of PancTu1 cells we added recombinant IL1RA, a natural IL1 receptor antagonist, and found that MSC-produced CCL2 as well as IL6 were significantly reduced (Figure 5A). Moreover, IL1RA was also able to block the induction of CCL2 and IL6 in MSCs by three additional pancreatic cell lines (Figure 5B) pointing to a general role of pancreatic cancer cell-derived IL1. Additionally, ELISA analyses revealed that both IL1α and IL1β are produced by pancreatic cancer cells (Figure 5C). As IL1 is a strong inducer of NF-κB we wondered whether PancTu1 supernatants would give rise to NF-κB activation in MSCs. Indeed, we found IκBα to be degraded in MSCs 5-30 min after exposure to PancTu1 supernatants, and NF-κB to be activated six-fold (Figure 5D). The NF-κB activation could be blocked by adding IL1RA to the pancreatic cancer cell supernatants (Figure 5D). In addition, an NF-κB inhibitor could stop the induction of CCL2 and IL6 by PancTu1 supernatants (Figure 5E) demonstrating the existence of an IL1-NF-κB-CCL2/IL6 pathway in MSCs in response to pancreatic cancer cells.

**Figure 5.**
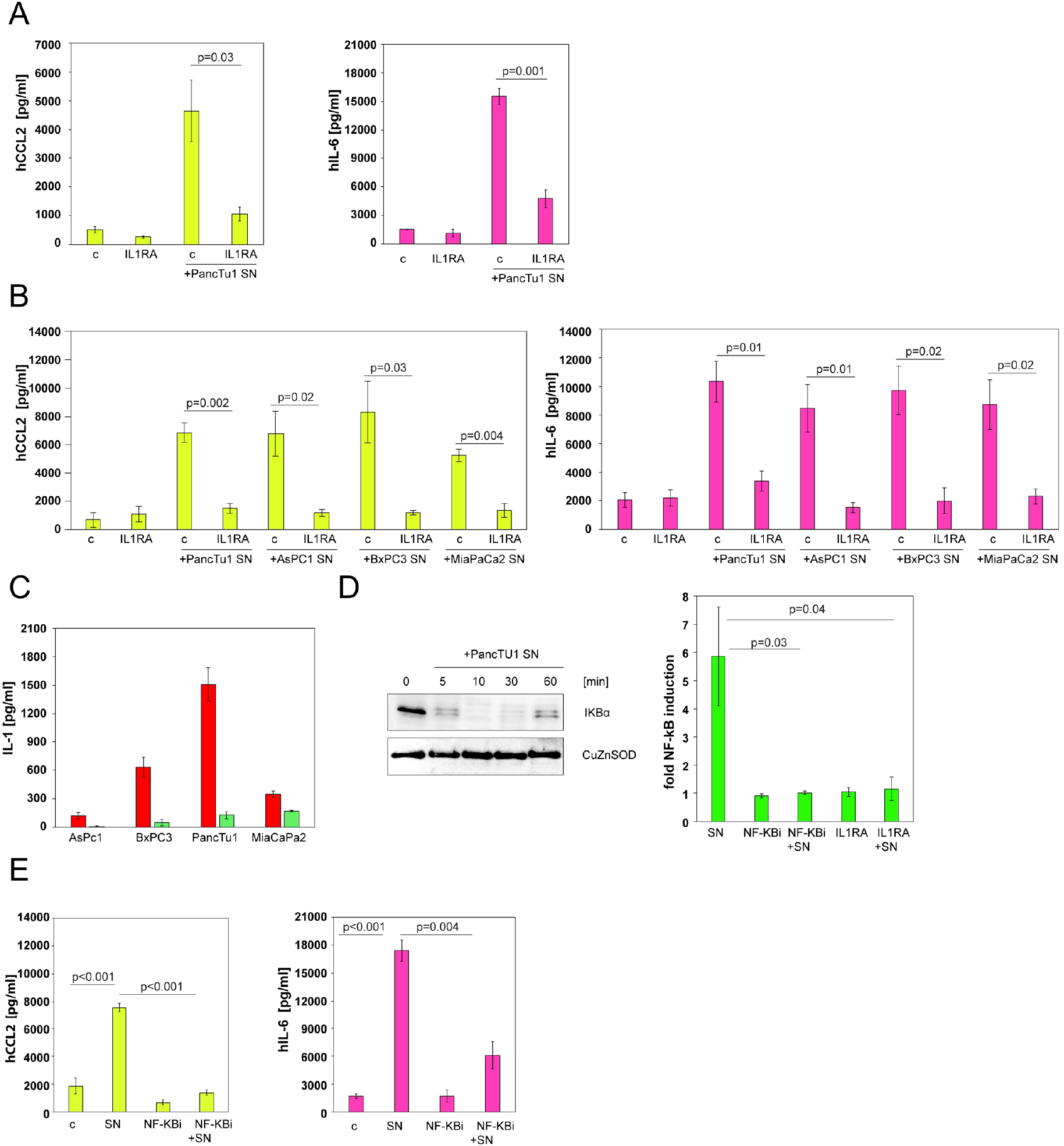
Pancreatic cancer-derived IL1 induces CCL2 and IL6 via NF-κB activation in MSCs. **A**. CCL2 (left) and IL6 (right) ELISAs of supernatants from human MSCs treated with conditioned medium (SN) from PancTu1 cells in the presence of IL1RA Carrier controls (c) and MSCs treated with IL1RA alone are also depicted in the graphs. Results are the mean ± SEM; n=3. **B**. CCL2 (left) and IL6 (right) ELISAs of supernatants from human MSCs treated with conditioned medium (SN) from different pancreatic cancer cell lines (PancTu1, AsPC1, BxPC3, MiaPaCa) in the presence of IL1RA Carrier controls (c) are also shown in the graphs. Results are the mean ± SEM; n=3. **C**. IL1α (red) and IL1β (green) ELISAs of supernatants from different pancreatic cancer cell lines (AsPC1, BxPC3, PancTu1, MiaPaCa) Results are the mean ± SEM; n=4. **D**. Western blot (left) of IκBα of human MSCs treated with conditioned medium from PancTu1 cells for different time points (0, 5, 10, 30, 60 min). The CuZnSOD blot serves as loading control. NF-κB reporter assay results (right) of MSCs treated with conditioned medium from PancTu1 cells (SN) as well as MSCs pre-treated with an NF-κB inhibitor (NF-κBi) and IL1RA, respectively. MSCs treated with NF-κB inhibitor or IL1RA alone were used as controls. Results are the mean ± SEM; n=4. **E**. CCL2 (left) and IL6 (right) ELISAs of supernatants from human MSCs treated with conditioned medium from PancTu1 cells (SN) with and without NF-κB inhibitor (NF-κBi). MSCs treated with carrier (c) or NF-κB inhibitor alone are shown as controls. Results are the mean ± SEM; n=3.

### IL1 gives rise to sustained CCL2 and IL6 production in MSCs and monocytic cells inhibit the induction of the two cytokines

Next, we analysed how sustained the CCL2 and IL6 production by MSCs was. For this we removed the supernatants from induced MSCs every three days and replaced it with fresh growth medium. At the end of the experiment we could still detect significantly elevated concentrations of CCL2 and IL6, albeit at lower levels than at earlier time points or when left to accumulate over nine days without medium change (Figure 6A). These results show that MSCs, once triggered, can persistently produce pro-metastatic cytokines, thereby achieving a potentially wide and long-lasting impact on surrounding cells.

**Figure 6.**
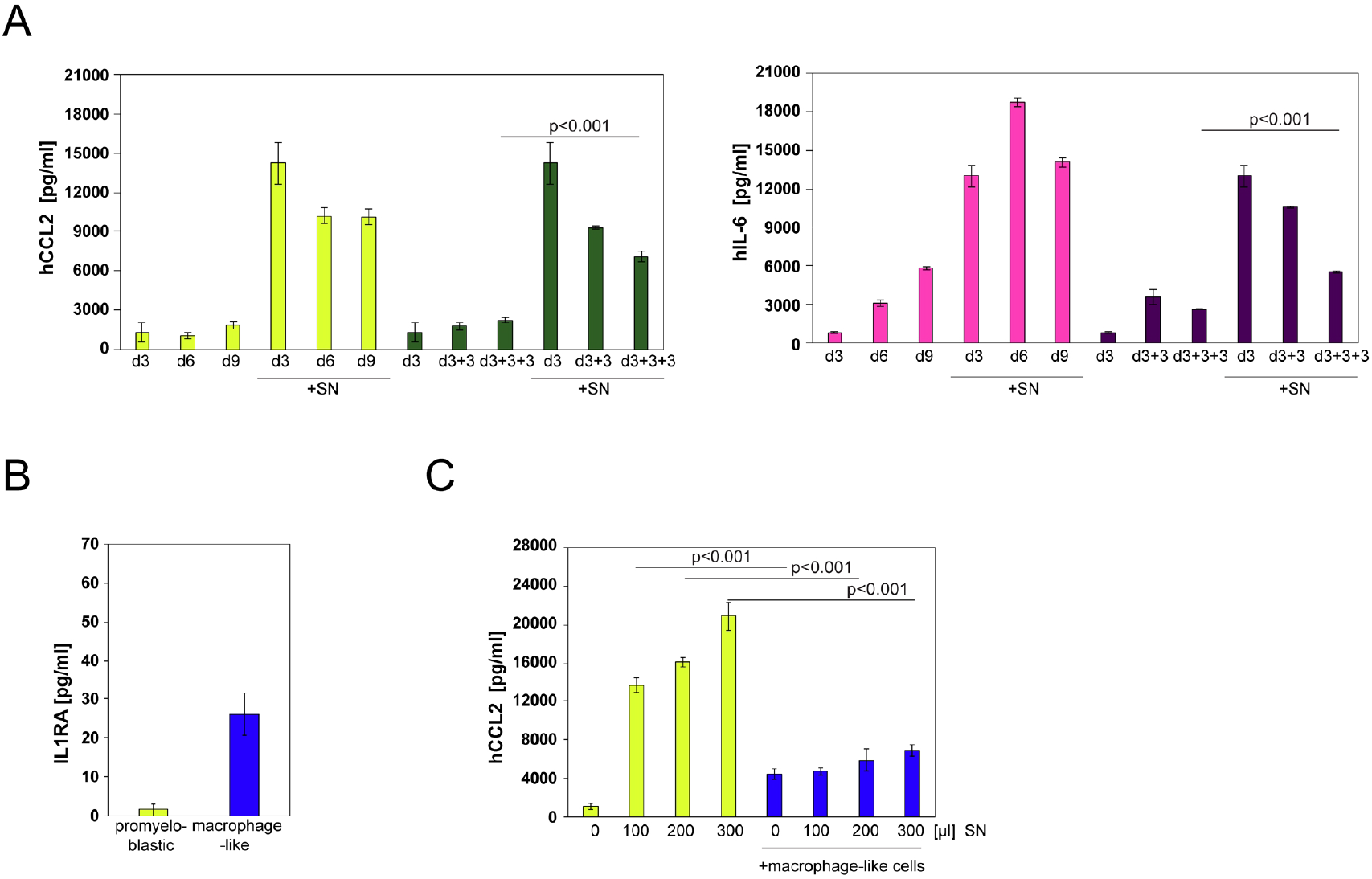
IL1 gives rise to sustained CCL2 and IL6 production in MSCs and monocytic cells inhibit the induction of the two cytokines. **A**. CCL2 (left) and IL6 (right) ELISAs of supernatants from human MSCs treated with conditioned medium from PancTu1 cells (SN) for three days followed by medium change every three days (dark green/purple bars). In comparison MSCs treated with conditioned medium from PancTu1 cells (SN) without medium change are shown (yellow/pink bars). MSCs without addition of conditioned medium serve as controls. Results are the mean ± SEM; n=3. **B**. IL1RA ELISA of supernatants of pro-myeloblastic and differentiated, macrophage-like cells. Results are the mean ± SEM; n=3. **C**. CCL2 ELISA of supernatants from human MSCs treated with different amounts of conditioned medium from PancTu1 cells (SN) in the presence of macrophage-like cells (blue bars). Supernatants from MSCs treated with different amounts of conditioned medium without macrophage-like cells are depicted as controls (yellow bars). Results are the mean ± SEM; n=3.

Anti-cancer therapeutic strategies targeting CCL2 activities have so far been tried rather unsuccessfully in clinical studies (Lim *et al*, 2016), which is contrary to robust anti-tumour responses in several pre-clinical cancer models (Chun *et al*, 2015; Fujimoto *et al*, 2009; Loberg *et al*, 2007; Qian *et al*, 2011; Sanford *et al*, 2013; Teng *et al*, 2017). We wondered whether our results could be linked to these problems, in particular the clinical finding that blocking CCL2 only transiently reduced serum CCL2 levels before they started to gradually come back and finally exceeded pretreatment baseline values (Sandhu *et al*, 2013). In this context, we found that macrophage-like monocytic cells express IL1RA (Figure 6B) and when we added such cells to the mix of PancTu1 cells and MSCs we could measure a significant reduction in CCL2 production (Figure 6C). Thus, CCL-2 recruited monocytic cells might form a negative feedback loop that limits the production of CCL2 in MSCs, and treatment with anti-CCL2 antibodies breaks this mechanism. Given these difficulties with CCL2-blocking approaches, we sought to investigate the Fas/FasL system in MSCs as a potential alternative target to block metastasis.

### Fas-deficient MSCs do not promote metastasis development

To further examine the importance of FasL signalling in MSC-promoted tumour progression we used MSCs established from MRL/MpJ-Fas^lpr^ mice (LPR-MSCs). These mice carry a mutation in the *Fas* gene and consequently LPR-MSCs show substantially reduced Fas protein expression as assessed by western blot and undetectable Fas surface expression measured by flow cytometry (Figure 7A). Consequently, in contrast to normal MSCs they did not respond to low-level FasL stimulation with increased growth demonstrating that they possess no functionally relevant expression of Fas (Figure 7B). Importantly, in animals bearing PancTu1 tumours the total number of LPR-MSC was lower when compared to mice injected with regular mMSCs, suggesting that the lack of Fas prevented proliferation of LPR-MSCs *in vivo* (Figure 7C). When we evaluated these LPR-MSC-injected mice for metastatic lesions we found none, as compared to 3 out of 4 animals in a control cohort injected with normal mMSCs (Figure 7D). These results indicate that without the FasL/Fas induced proliferation, the static number of LPR-MSCs do not produce sufficient levels of pro-metastatic factors to induce metastasis. Thus, targeting Fas signalling in MSCs might be a strategy to prevent tumour cells from progressing and metastasing, and in addition, knocking-out Fas by gene editing might be an approach to make MSCs safer for cell therapeutic approaches.

**Figure 7.**
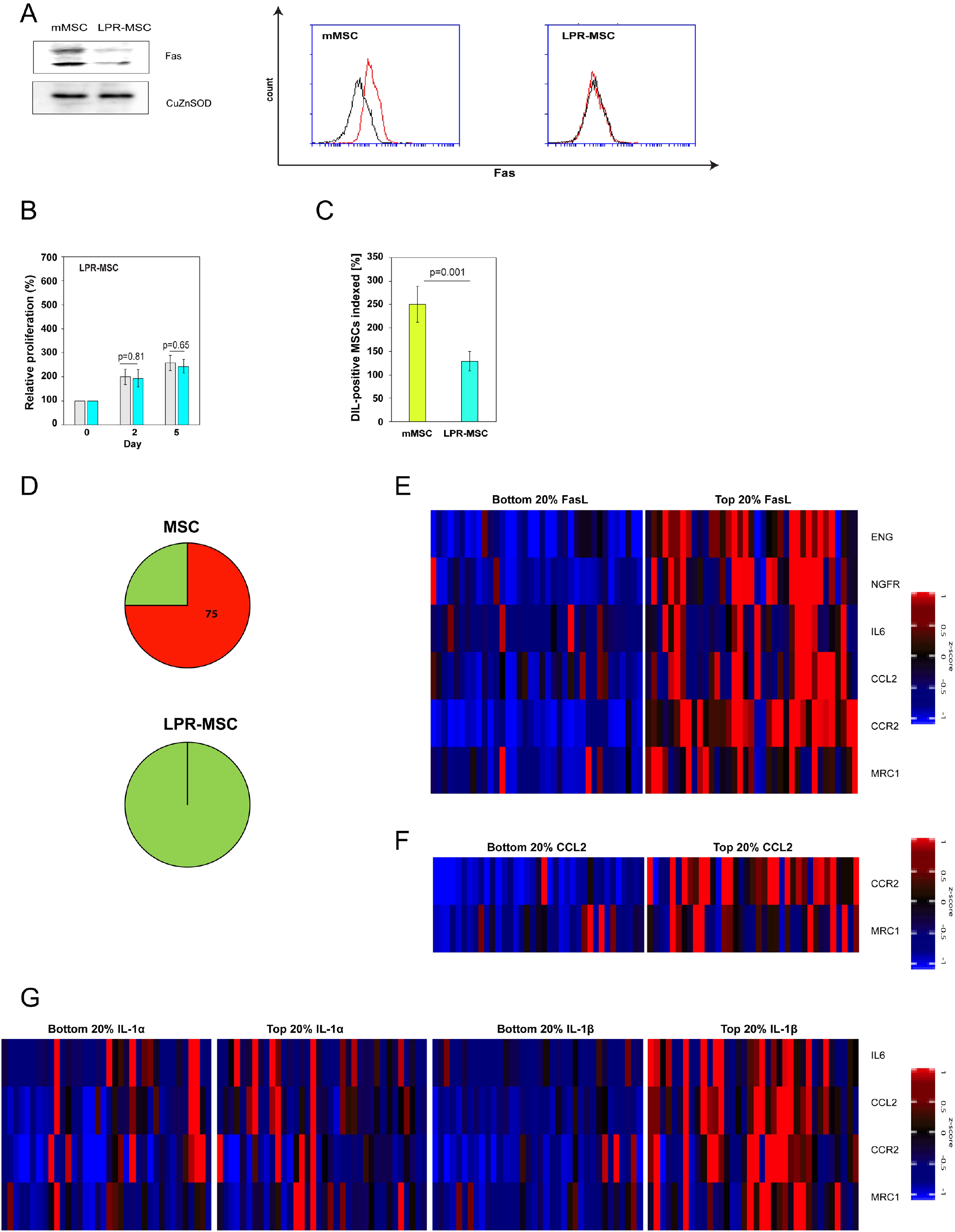
Fas-deficient MSCs do not promote metastasis development. **A**. Western blot (left) for Fas expression in murine MSCs (mMSCs) and LPR-MSCs. The CuZnSOD blot serves as loading control. Flow cytometric analysis (right) of Fas surface expression (red) on murine MSCs (mMSCs) and LPR-MSCs. The isotype control is shown in black. **B**. LPR-MSCs were treated with 0.2 ng/ml of FasL for 2 and 5 days and cell numbers determined (turquoise). MSCs treated with carrier are shown as controls (grey). The numbers of LPR-MSCs at the start of FasL treatment (Day 0) were set to 100%. Results are the mean ± SEM; n=6. **C**. Relative numbers of murine MSCs (mMSCs) and LPR-MSCs found in tumour-burdened (PancTu1) animals after 28 days. Each cohort consisted of four animals. **D**. Incidence of lung metastases in PancTu1 tumour-burdened animals after systemic administration of murine MSCs (mMSCs) or LPR-MSCs. Each cohort consisted of four animals. **E**. Heatmap of MSC-markers (ENG, NGFR), cytokines (IL6, CCL2) and monocytic markers (CCR2, MRC1) that are significantly co-expressed (p≤0.001) with FasL. The heatmap shows 20% of samples with highest and lowest FasL expression, respectively. n=186. **F**. Heatmap of monocytic markers (CCR2, MRC1) significantly co-expressed (p≤0.001) with CCL2 showing 20% of samples with highest and lowest CCL2 expression, respectively. n=186. **G**. Heatmap of cytokines (IL6, CCL2) and monocytic markers (CCR2, MRC1) that are co-expressed with IL1α (left) and IL1β (right; p≤0.001) showing 20% of samples with highest and lowest IL1α/IL1β expression, respectively. n=186.

Our results establish a network of cancer cells and MSCs responding to and acting via FasL, IL1, CCL2 and IL6. To evaluate whether this FasL-IL1-CCL2-IL6 circuit exists in pancreatic cancer patients, we subjected RNA sequencing (RNA-seq) data from a cohort of 186 pancreatic adenocarcinoma samples to bioinformatic co-expression analysis. It revealed that FasL expression showed a remarkable and significant positive correlation with the expression of two MSC markers (NGFR and Endoglin), CCL2 and IL6 as well as monocyte specific markers (CCR2 and MRC1) (Figure 7E and Supplementary Figure 3A). Furthermore, high CCL2 expression was associated with increased levels of the monocytic markers CCR2 and MRC1 (Figure 7F and Supplementary Figure 3B). Finally, IL1β, but not IL1α, was co-expressed with CCL2, IL6, CCR2 and MRC1 (Figure 7G and Supplementary Figure 3C). In contrast to FasL, TRAIL (TNFSF10) expression was not correlated to the two MSC markers NGFR and Endoglin (Supplementary Figure 3D). These results demonstrate that in human pancreatic cancer FasL appears to be able to give rise to an increase in intra-tumoural MSCs that in response to IL1 prompt the production of CCL2 and IL6. These cytokines, aside from having direct effects on cancer cells, also appear to mediate the recruitment of monocytic cells (Figure 8).

**Figure 8.**
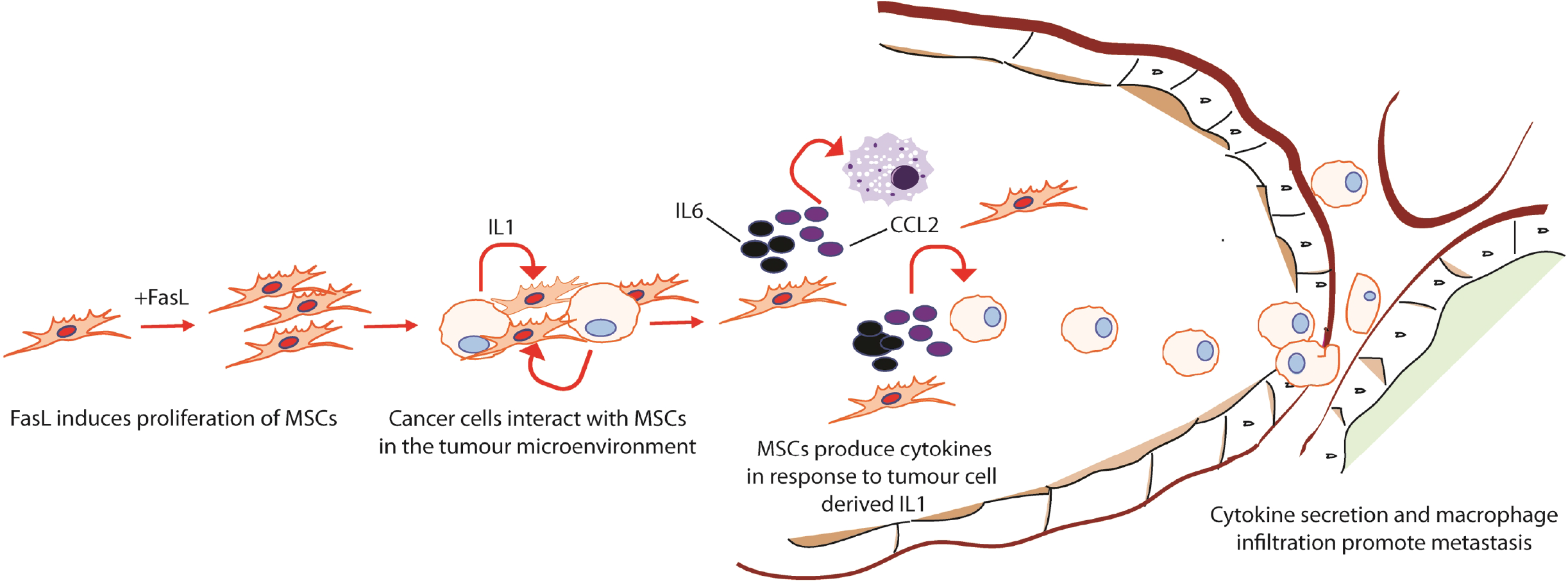
Schematic diagram of FasL-induced mechanism. Model of the effect of FasL on MSCs and their role in the crosstalk with pancreatic cancer cells in the TME.

## Discussion

MSCs have been shown to promote tumour metastasis in various cancer models (Albarenque *et al*., 2011; Jung *et al*, 2013; Karnoub *et al*., 2007; Luo *et al*, 2014; Xu *et al*, 2009), but not all underlying cellular and molecular mechanisms have been clarified. Here, we show that Fas signalling in MSCs is required for their pro-metastatic activity, as MSCs lacking Fas expression do not lead to tumour progression in a pancreatic cancer model. Mechanistically, we found that MSCs are resistant to FasL-induced apoptosis and that low concentrations of FasL triggered so called Fas-threshold signalling promoting proliferation. This response was absent in LPR-MSCs. In line with these *in vitro* data we found fewer LPR-MSCs as compared to regular MSCs in tumour-burdened mice. These results link FasL/Fas-mediated-proliferation of MSCs to their capacity to promote metastasis development. While tumour-promoting activities of Fas and its ligand FasL have been reported before (Peter *et al*., 2015), our findings expand this role to MSCs in the TME.

There are several different possible sources of FasL in the context of a growing tumour. One possibility is that tumour-associated endothelial cells express FasL on their surface providing the proliferation signal to MSCs when they infiltrate the tumour tissue (Le Gallo *et al*, 2017; Malleter *et al*., 2013; Motz *et al*, 2014). Other reports have indicated that (pancreatic) cancer cells themselves express FasL (Ungefroren *et al*, 1998), but we were unable to confirm these findings. Regardless of the origin of FasL, we found that in human pancreatic cancer, enhanced FasL expression is associated with higher levels of MSC-markers in the tumour. The FasL-induced larger number of MSCs consequently provide increased capacity to produce pro-metastatic factors.

In a breast carcinoma model CCL5 was identified as the pro-metastatic cytokine that causes MSC-induced metastasis development (Karnoub *et al*., 2007). However, we were unable to find elevated CCL5 when MSCs were mixed with pancreatic cancer cells. Instead, we discovered CCL2 and IL6 to be induced after interaction of pancreatic cancer cells with MSCs, identifying MSCs as a novel cellular production site for CCL2 and IL6 in the TME. Importantly, this induction gave rise to prolonged expression of the two cytokines providing a longer window of action in which they can exert their pro-metastatic activities. Furthermore, we discovered that pancreatic cancer cell-derived IL1 is responsible for the CCL2 and IL6 induction via NF-κB activation in MSCs, as both an NF-κB inhibitor and the natural antagonist of the IL1 receptor, IL1RA, could fully inhibit the CCL2 and IL6 induction.

The two cytokines, CCL2 and IL6, possess well documented pro-metastatic activities (Ara & Declerck, 2010; Lim *et al*., 2016). IL6 is a pleiotropic cytokine known to be involved in chronic inflammation and auto-immune diseases, but has also been shown to contribute to the generation of a pro-tumorigenic microenvironment regulating angiogenesis and metastasis (Garbers *et al*, 2018; Kang *et al*, 2019; Kumari *et al*, 2016; Murakami *et al*, 2019). In the serum of pancreatic cancer patients, it was found at elevated levels, and correlates with advanced tumour stage and poor survival (Miura *et al*, 2015). Furthermore, knocking-out IL6 in a transgenic mouse model of pancreatic cancer inhibited the maintenance and progression of pancreatic cancer precursor lesions (Zhang *et al*, 2013). As we could not measure a direct impact of IL6 on the invasiveness of pancreatic cancer cells, it is likely that IL6 exerts its pro-metastatic effect via indirect mechanisms engaging additional cell types (Kumari *et al*., 2016). CCL2 is a small chemokine that belongs to the CC chemokine family (Griffith *et al*, 2014). Although first described as a chemotactic molecule for monocytes, basophils, T-lymphocytes and NK cells with physiological roles in regulating inflammation, more recent studies revealed a pro-tumorigenic function for CCL2 favouring cancer development and subsequent metastasis (Chow & Luster, 2014; O’Connor *et al*, 2015). Indeed, cancer progression and poor prognosis have been linked to enhanced levels of CCL2 in a number of cancer types including pancreatic cancer (Kudo-Saito *et al*, 2013; Li *et al*, 2009; Lu *et al*, 2006; Sanford *et al*., 2013; Yoshidome *et al*, 2009). Therefore, CCL2 was proposed as a therapeutic target, which led to several clinical trials with anti-CCL2 antibodies in solid and metastatic cancers (Lim *et al*., 2016). However, it became clear in these studies that CCL2 could only be transiently suppressed, and over time the CCL2 concentration increased to levels beyond pre-treatment baseline values. Moreover, none of the treated patients showed an objective anti-tumour response (Pienta *et al*, 2013; Sandhu *et al*., 2013; Vela *et al*, 2015). It is known that CCL2 can exert direct actions on cancer cells, as we demonstrated, as well as initiate the infiltration of pro-metastatic cell types including monocytes (Hartwig *et al*, 2017; Qian *et al*., 2011). We hypothesised that monocytic cells might also be involved in the regulation of CCL2 expression in MSCs, as we and others have shown that they express IL1RA (Arend *et al*, 1985; Arend *et al*, 1990). Indeed, when we mixed monocytic cells with MSCs and conditioned medium from pancreatic cancer cells, CCL2 production was no longer induced. These findings provide a possible explanation for the rebounding CCL2 levels observed in clinical tests of anti-CCL2 antibodies. Despite failing in clinical tests, the anti-CCL2 antibodies afforded an initial reduction in the number of monocytic cells in the tumours. However, as monocytic cells normally carry IL1RA to tumours they can limit the production of CCL2. Thus, while blocking CCL2 will give rise to a reduction in monocytic cells in the tumour, it will also lead to continuous and unchecked CCL2 production by MSCs. The loss of this homeostatic feedback mechanism can eventually overwhelm the capacity of the neutralising antibodies leading to an overshoot of the CCL2 levels as observed in clinical studies. Thus, targeting factors such as FasL or IL1, that act on MSCs, might offer better alternatives.

Overall, our results establish MSCs as a central cellular player in the TME of pancreatic cancer orchestrating a string of molecular and cellular changes that contribute to tumour progression and metastasis development.

## Acknowledgments

We thank Stella-Maris Albarenque and Jamie Moore for their assistance with this project.

## Author contribution

AM, VBT, GNB and RMZ and supervised the study; AM performed animal experiments and in vitro experiments, and analysed data; CT and GNB completed and analysed *in vitro* studies; CTC analysed patient data and generated heatmaps; AM and RMZ wrote the paper; VBT, GNB and CTC edited the manuscript.

