## Supplemental info for "Fas-threshold signalling in MSCs causes tumour progression and metastasis"

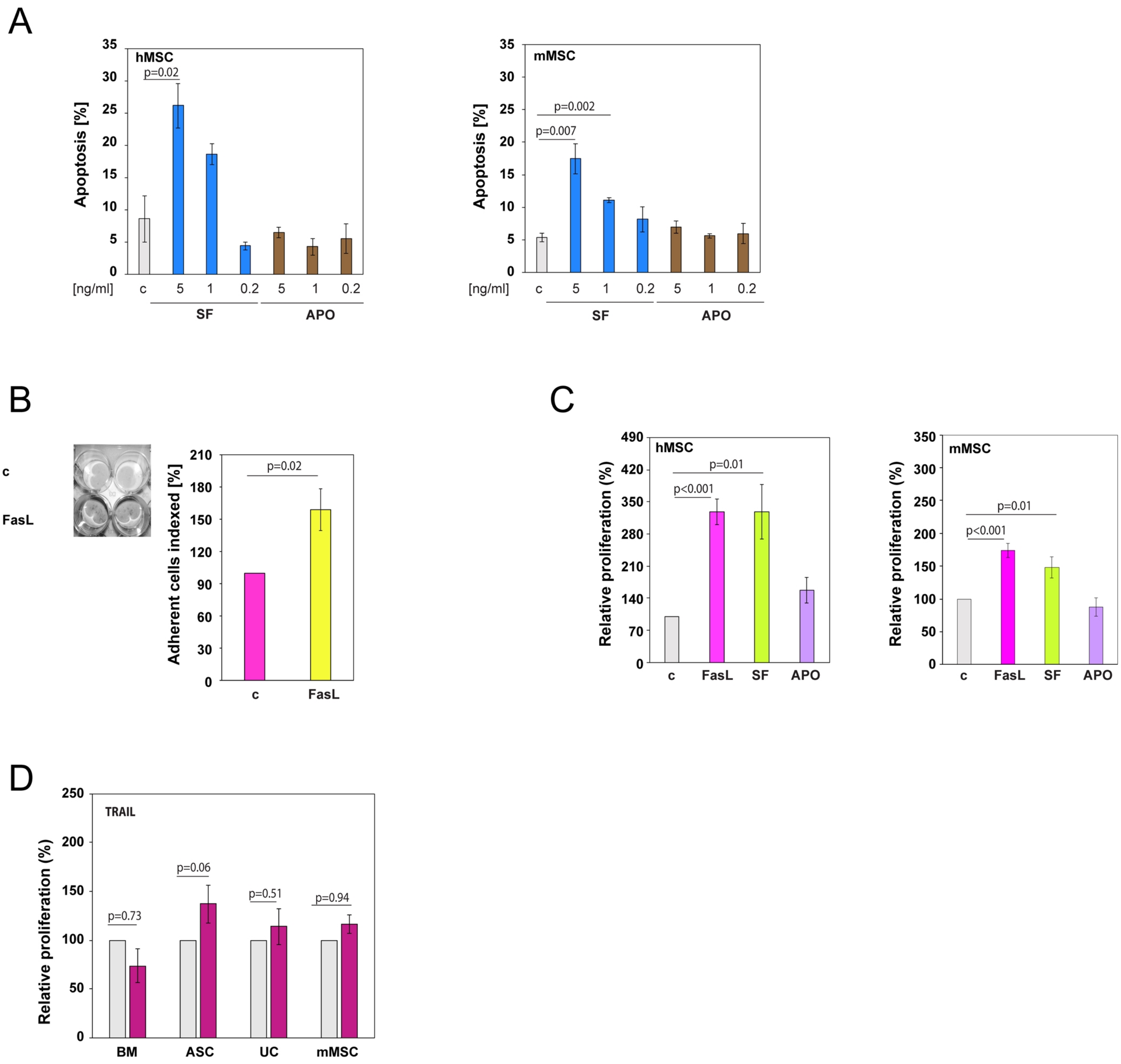
A
hMSC
Apoptosis [%]
35
30
25
20
15
10
5
0
p=0.02
[ng/ml]
c
5
1
0.2
5
1
0.2
SF
APO
mMSC
Apoptosis [%]
p=0.002
p=0.007
[ng/ml]
SF
APO
B
c
FasL
Adherent cells indexed [%]
210
180
150
120
90
60
30
0
p=0.02
c
FasL
C
hMSC
Relative proliferation (%)
490
420
350
280
210
140
70
0
p=0.01
p<0.001
c
FasL
SF
APO
mMSC
Relative proliferation (%)
350
300
250
200
150
100
50
0
p=0.01
p<0.001
c
FasL
SF
APO
D
TRAIL
Relative proliferation (%)
250
200
150
100
50
0
p=0.73
p=0.06
p=0.51
p=0.94
BM
ASC
UC
mMSC

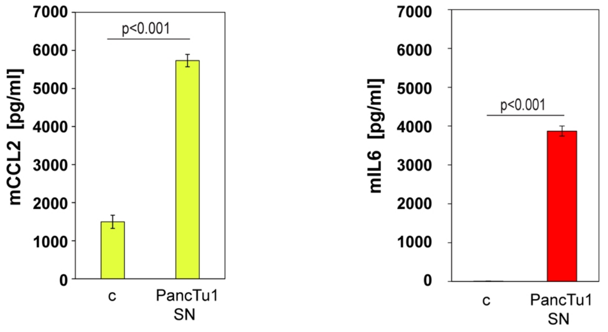

mCCL2 [pg/ml]
7000
6000
5000
4000
3000
2000
1000
0
p<0.001
c
PancTu1
SN
mIL6 [pg/ml]
7000
6000
5000
4000
3000
2000
1000
0
p<0.001
c
PancTu1
SN

A

| Queried Gene | Correlated Gene | Cytoband | Spearman's Correlation | p-Value | q-Value |
|---|---|---|---|---|---|
| FasL | ENG | 9q34.11 | 0.486892107 | 6.35E-12 | 1.46E-10 |
| FasL | NGFR | 17q21.33 | 0.477255415 | 1.87E-11 | 4.02E-10 |
| FasL | IL6 | 7p15.3 | 0.357856079 | 1.00756E-06 | 7.8E-06 |
| FasL | CCL2 | 17q12 | 0.464746975 | 7.20E-11 | 1.37E-09 |
| FasL | CCR2 | 3p21.31 | 0.754873531 | 6.97E-34 | 4.80E-31 |
| FasL | MRC1 | 10p12.33 | 0.388787417 | 8.91E-08 | 8.88E-07 |

B

| Queried Gene | Correlated Gene | Cytoband | Spearman's Correlation | p-Value | q-Value |
|---|---|---|---|---|---|
| CCL2 | CCR2 | 3p21.31 | 0.551671976 | 1.75E-15 | 2.82E-13 |
| CCL2 | MRC1 | 10p12.33 | 0.299771905 | 5.04422E-05 | 0.00031 |

C

| Queried Gene | Correlated Gene | Cytoband | Spearman's Correlation | p-Value | q-Value |
|---|---|---|---|---|---|
| IL1α | IL6 | 7p15.3 | 0.112352134 | 0.136520159 | 0.28821 |
| IL1α | CCL2 | 17q12 | 0.188513095 | 0.011977552 | 0.05288 |
| IL1α | CCR2 | 3p21.31 | 0.039296736 | 0.603547706 | 0.75058 |
| IL1α | MRC1 | 10p12.33 | 0.050301467 | 0.506116353 | 0.67539 |

| Queried Gene | Correlated Gene | Cytoband | Spearman's Correlation | p-Value | q-Value |
|---|---|---|---|---|---|
| IL1β | IL6 | 7p15.3 | 0.408456831 | 1.66E-08 | 6.6E-06 |
| IL1β | CCL2 | 17q12 | 0.508957168 | 4.72E-13 | 9.42E-09 |
| IL1β | CCR2 | 3p21.31 | 0.430545697 | 2.21E-09 | 1.8E-06 |
| IL1β | MRC1 | 10p12.33 | 0.303565555 | 4.00117E-05 | 0.00143 |

D

| Queried Gene | Correlated Gene | Cytoband | Spearman's Correlation | p-Value | q-Value |
|---|---|---|---|---|---|
| TNFSF10 | ENG | 9q34.11 | 0.134123803 | 0.07510725 | 0.2103 |
| TNFSF10 | NGFR | 17q21.33 | 0.087983241 | 0.244218542 | 0.44405 |

**Supplementary Figure 1**

**A.** Apoptosis measurements in mMSCs (left) and hMSCs (right) after treatment with multimeric FasL (SF) and anti-APO-1-3 (APO) for 24 h, at concentrations ranging from 0.2-5 ng/ml. Controls (c) were treated with carrier only. Results are the mean ± SEM; n=3.

**B.** Growth of hMSCs following treatment with carrier (c) or 0.2 ng/ml of FasL for 5 days. The cells were visualised by crystal violet stain. A representative image is shown on the left and quantification of the results on the right. Growth of the untreated cells after 5 days was set to 100%. Results are the mean ± SEM; n=5.

**C.** hMSC (left) and mMSCs (right) were treated with FasL, multimeric FasL (SF) and anti-APO-1-3 (APO) as shown in the figure for 2 days. The numbers of MSCs in the carrier controls (c) were set to 100%. Results are the mean ± SEM; n=4 (hMSC); n=8 (mMSC).

**D.** Human MSCs from bone marrow (BM), adipose tissue (ASC) and umbilical cord (UC) as well as murine MSCs (mMSC) were treated with 0.2 ng/ml of TRAIL for 2 days and cell numbers determined (purple). MSCs treated with carrier are shown as controls (grey). The numbers of MSCs at the start of TRAIL treatment were set to 100%. Results are the mean ± SEM; (n=4)

**Supplementary Figure 2**

CCL2 (left) and IL6 (right) ELISAs of supernatants from mMSCs treated with conditioned medium from PancTu1 cells (SN) for three days. MSCs treated with unconditioned medium served as controls (c). Results are the mean ± SEM; n=3.

**Supplementary Figure 3**

**A.** Correlation coefficients and p-values of co-expression relationships of FasL with the listed genes.

**B.** Correlation coefficients and p-values of co-expression relationships of CCL2 with the listed genes.

**C.** Correlation coefficients and p-values of co-expression relationships of IL1$\alpha$ (top) and IL1$\beta$ (bottom) with the listed genes.

**D.** Correlation coefficients and p-values of co-expression relationships of TNFSF10 (TRAIL) with the listed genes.
